# A re-analysis of the data in Sharkey et al.’s (2021) minimalist revision reveals that BINs do not deserve names, but BOLD Systems needs a stronger commitment to open science

**DOI:** 10.1101/2021.04.28.441626

**Authors:** Rudolf Meier, Bonnie B. Blaimer, Eliana Buenaventura, Emily Hartop, Thomas von Rintelen, Amrita Srivathsan, Darren Yeo

## Abstract

Halting biodiversity decline is one of the most critical challenges for humanity, but monitoring biodiversity is hampered by taxonomic impediments. One impediment is the large number of undescribed species (here called “dark taxon impediment”) while another is caused by the large number of superficial species descriptions which can only be resolved by consulting type specimens (“superficial description impediment”). Recently, Sharkey et al. (2021) proposed to address the dark taxon impediment for Costa Rican braconid wasps by describing 403 species based on barcode clusters (“BINs”) computed by BOLD Systems. More than 99% of the BINs (387 of 390) are converted into species by assigning binominal names (e.g., BIN “BOLD:ACM9419” becomes *Bracon federicomatarritai*) and adding a minimal diagnosis (usually consisting only of a consensus barcode). We here show that many of Sharkey et al.’s species are unstable when the underlying data are analyzed using different species delimitation algorithms. Add the insufficiently informative diagnoses, and many of these species will become the next “superficial description impediment” for braconid taxonomy because they will have to be tested and redescribed after obtaining sufficient evidence for confidently delimiting species. We furthermore show that Sharkey et al.’s approach of using consensus barcodes as diagnoses is not functional because it cannot be consistently applied. Lastly, we reiterate that COI alone is not suitable for delimiting and describing species and voice concerns over Sharkey et al.’s uncritical use of BINs because they are calculated by a proprietary algorithm (RESL) that uses a mixture of public and private data. We urge authors, reviewers, and editors to maintain high standards in taxonomy by only publishing new species that are rigorously delimited with open-access tools and supported by publicly available evidence.

---

After being principally an academic subject for many decades, biodiversity declines are now also a major topic for decision makers (e.g., economists, CEOs, political leaders) (World Economic Forum, 2020). This heightened interest has highlighted once more that one of the most important tasks in science is unfinished: the discovery and description of earth’s biodiversity (May, 2011). Most of the species-level diversity and a significant proportion of biomass are concentrated in taxonomically poorly known taxa that are nowadays often referred to as “dark taxa” (e.g., bacteria, fungi, invertebrates) (Hausmann et al., 2020). Such groups are usually avoided by taxonomists because they combine high diversity with high abundance. Yet, dark taxa are a major cause for the “taxonomic impediment” that is the result of a range of issues. One is the large number of undescribed species (here called “dark taxon impediment”) while another is species descriptions that are too superficial by today’s standards to allow for the identification of the species without inspecting type specimens, collecting additional data, and re-describing the species (here called “superficial description impediment”). Addressing the superficial description impediment is particularly time-consuming because it requires museum visits/loans, type digitization, and possibly lengthy searches for misplaced specimens. Eventually, the species have to be re-described, making the total process of getting to a useful species description take twice the amount of time needed if there had not been a poor-quality legacy description. The reality is that new species descriptions could be accelerated manifold if taxonomists did not have to spend so much time on improving existing descriptions.

What the 21^st^ century undoubtedly needs is more species descriptions, but what should be avoided is creating new “superficial description impediments” in the form of large numbers of poorly supported and described species. Yet, this is what Sharkey et al. (2021) propose when they state: “… we view barcode-based descriptions as a first pass in an iterative approach to solve the taxonomic impediment of megadiverse and under-taxonomically resourced groups that standard technical and biopolitical approaches have not been able to tackle.” Sharkey et al. delegate the critical “iterative” work to future generations of taxonomists. They will have to start their own revisions by first revisiting the species descriptions, types, and specimens of Sharkey et al.’s (2021) species to resolve the species boundaries based on data that should have been collected and analyzed at the time of description.

Upon closer inspection, Sharkey et al.’s “minimalist revision” relies on three core assumptions/techniques which are here shown to be problematic. The first is that COI barcodes are sufficiently decisive and informative that they can be used as the only/main data source for delimiting and describing species. The second is that species can be diagnosed unambigously using only a consensus barcode. The third is that it is sufficient to use a particular type of barcode cluster (“BIN”) that is delimited by a proprietary algorithm that is only installed on BOLD systems.

## (1) Are COI barcodes sufficiently informative for delimiting and describing species?

Sharkey et al.’s revision covers 390 BINs obtained from BOLD. Of these, >99% (387) are converted into species. This is surprising because virtually all studies that have looked into the stability of barcode clusters have found that the cluster composition varies depending on which barcode clustering algorithm is used (Virgilio et al., 2010; Kekkonen and Hebert, 2014; Srivathsan et al., 2019; Yeo et al., 2020; Hartop et al., 2021); i.e., barcode data is not sufficiently decisive that they can be described as species – rather, species delimitation must integrate multiple forms of data. We here first tested whether the barcode data used in Sharkey et al. (2021) is unusually decisive and could be used without validation from another source. Conventionally, such re-analyses of data from published studies are straightforward because the data are made available by the authors. However, the “molecular data” in the supplementary material of Sharkey et al. (2021) consist only of neighbor-joining (NJ) trees. We thus proceeded to obtain the sequences directly from BOLD by consulting the BIN numbers in the revision and NJ trees in the supplement. Note, however, that BINs are calculated based on public and private data with the ratio between the two apparently being approximately 3:1 (see below). In addition, the NJ trees in the supplement contained non-Costa Rican BINs from other Neotropical countries that were not covered by the revision. Consequently, we ended up with three datasets of increasing size: (1) public barcodes for only those BINs described by Sharkey et al. (2021), (2) public barcodes for all BINs found on the NJ trees in the supplementary materials, and (3) all public barcodes in the BOLD database for the 11 braconid subfamilies covered by the revision of the Costa Rican fauna (see supplementary data).

All three datasets were analyzed with several species delimitation methods as recommended by Carstens et al. (2013). The overall analyses followed Yeo et al. (2020): objective clustering with uncorrected p-distance thresholds at 1-5% using TaxonDNA (Meier et al., 2006), ASAP – a new implementation of ABGD – where we only kept the results obtained with the top five partitions (= lowest p-values) (Puillandre et al., 2012a; Puillandre et al., 2021), and Single-rate PTP (Zhang et al., 2013) based on maximum likelihood trees generated from the barcode data with RAxML v.8 (Stamatakis, 2014). These re-analyses were used to test whether the different methods unambiguously supported the BINs that were described as species by Sharkey et al. (2021). Arguably, this is a minimum requirement for using BIN=Species as default because it is hard to justify describing barcode clusters as species if the former are not consistent across methods.

We find that in dataset 1, 131 of the 401 BINs (32.7%) conflict with at least one of the species delimitation treatments. The instability of these BINs increases as more *COI* evidence is included (Fig. 1), with 283 of the 615 BINs (46.0%) from dataset 2, and 2912 of the 3896 BINs (74.7%) from dataset 3 in conflict with the results of at least one of the analyses. The corresponding numbers for only those 401 BINs described as species in Sharkey et al. (2021) are 138 (34.4%: dataset with all Neotropical barcodes from BOLD) and 276 (68.8% dataset with all braconid barcodes from BOLD). This means that as braconid barcodes are sampled more densely across the Neotropics and the world, an increasing number of the BINs described as species in Sharkey et al. (2021) contain specimens that are assigned to barcode clusters that disagree with the BIN assignment from BOLD. Overall, this reanalysis shows that COI barcodes contain ambiguous signal when it comes to delimiting many species, and that this problem increases with sampling.

**Figure 1.**
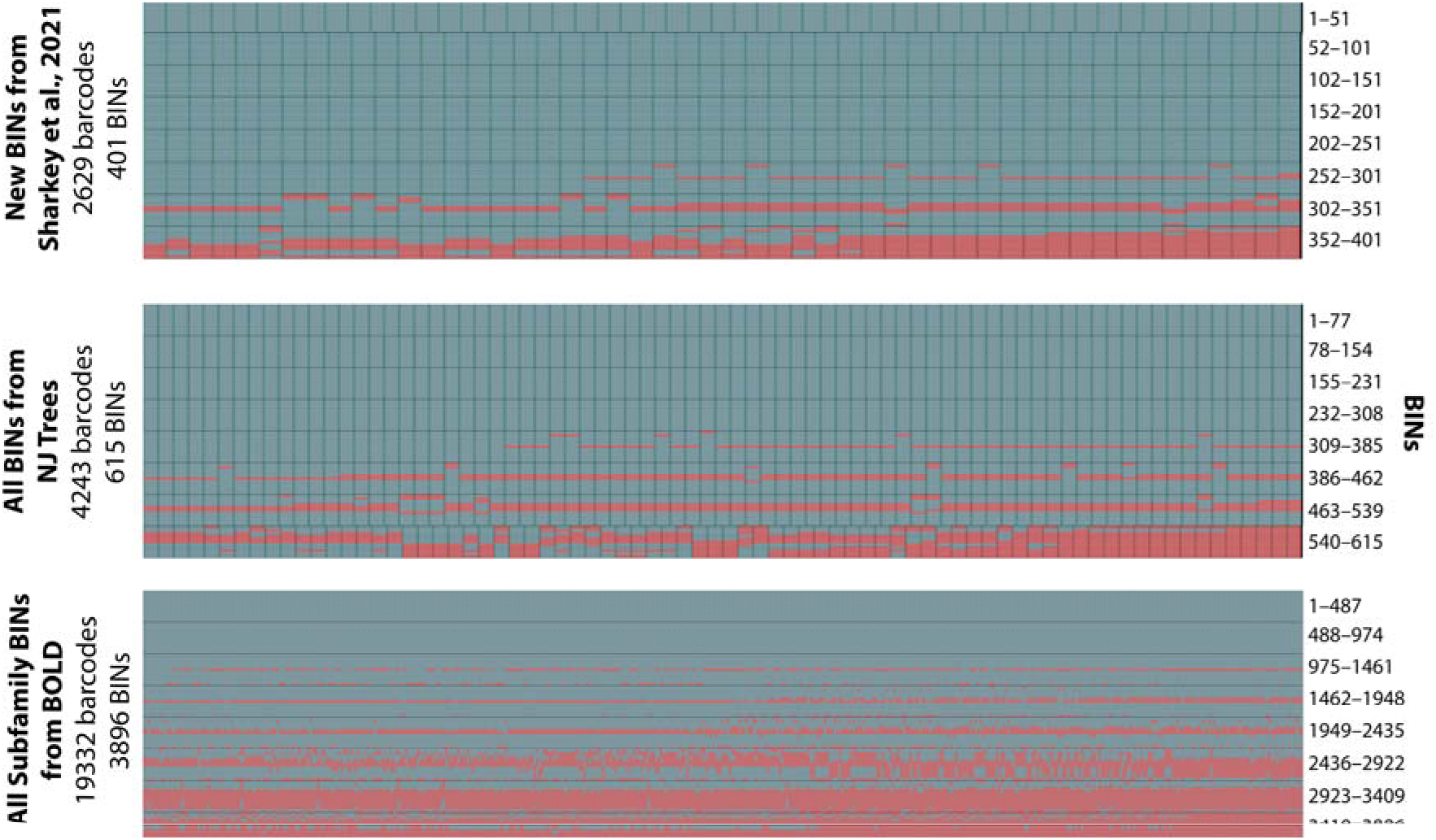
Heatmaps showing incongruence between species delimitations for three datasets of increasing size. In order to accommodate a large number of BINs, each heatmap is split into 8 blocks that are separated by dark horizontal lines (number of bins represented in each block is provided). Rows within each block represent species delimitation algorithms (PTP, objective clustering: 1 – 5% p-distance, ASAP: top 5 priors) and BINs in columns (blue = congruent BINs; pink = incongruent BIN).

To obtain lower-bound estimates for incongruence, we identified the parameters for each species delimitation algorithm (objective clustering, ASAP) whose application yielded the largest number of barcode clusters that are identical to BINs; i.e., the parameters from each algorithm that maximized congruence with BINs. We find that 38 of the 401 BINs (9.48%) described in the paper are incongruent even after the methods were optimized to replicate BINs as well as possible (Fig. 2). Incongruence again increases as more of the available barcodes and BINs are included [dataset 2: 82 of the 615 BINs incongruent (13.33%), dataset 3: 1340 of the 3896 BINs (34.39%) incongruent]. These analyses illustrate that the available *COI* data provide ambiguous support for 10-30% of the BINs described as new species in Sharkey et al. (2021). This finding is congruent with virtually all studies that have investigated the congruence between barcode clusters obtained with different clustering methods (Virgilio et al., 2010; Kekkonen and Hebert, 2014; Srivathsan et al., 2019; Yeo et al., 2020; Hartop et al., 2021). Furthermore, there are good reasons to assume that 10-30% is an underestimate of the true number of clusters that require other forms of data for resolving species boundaries because denser *COI* sampling will close barcoding gaps between species, thus leading to further uncertainty with regard to the boundaries of barcode clusters (Bergsten et al., 2012).

**Figure 2.**
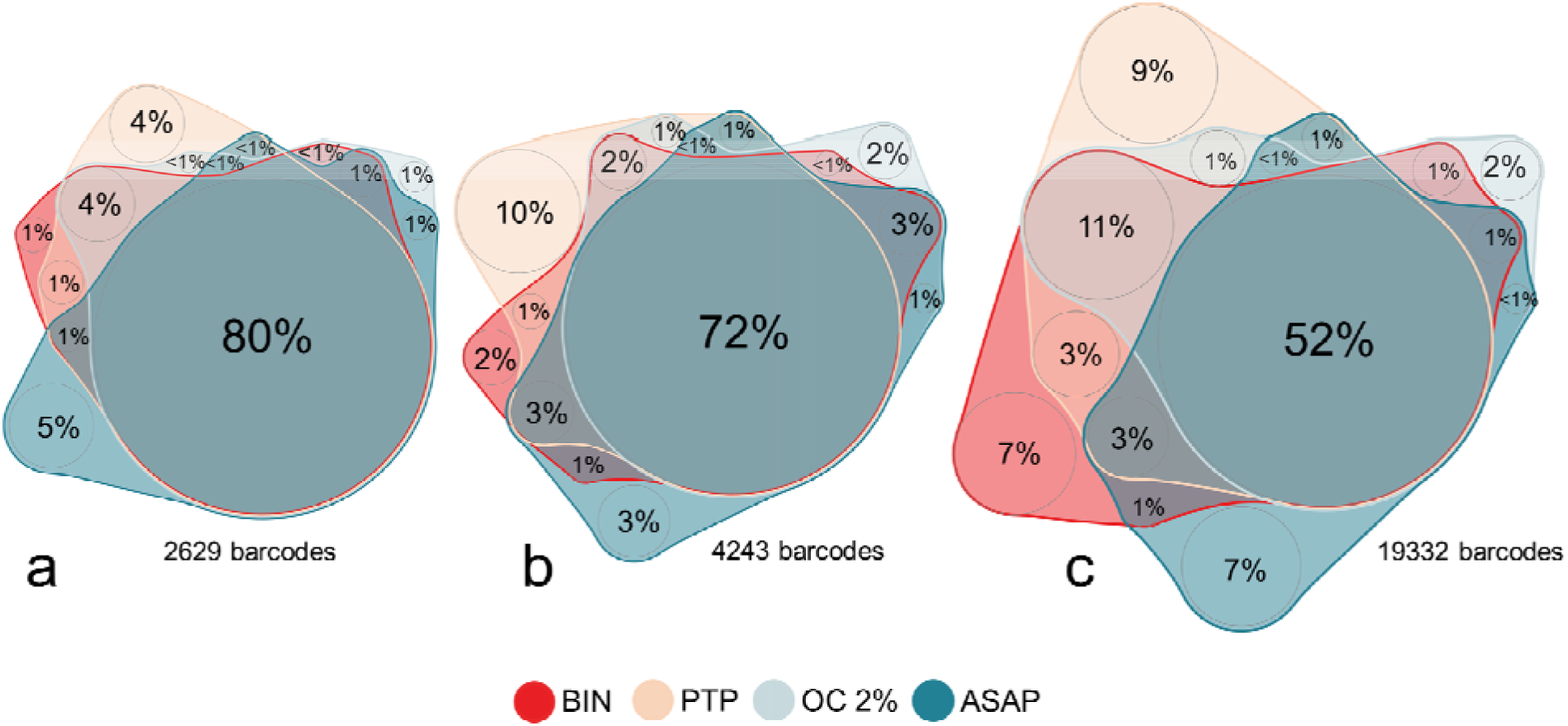
Barcode cluster congruence across methods using optimal thresholds for BIN, PTP, Objective Clustering (OC) (2%) and ASAP (d −3) for (a) dataset I, (b) dataset II, (c) dataset III.

Sharkey et al. (2021) are aware that broader geographical sampling regularly leads to BINs that contain multiple species: “Braconid specimens from the following New World countries appear to be relatively well-sampled in BOLD: Canada, USA, Belize, Argentina, French Guiana, and Mexico. There is a small number of cases where specimens from these countries fall in the same BIN as one of our Costa Rican species, but they were not studied. More sampling between these disparate localities, and more genomic and/or morphological and behavioral data will help resolve these species-level cases, which are beyond the scope of this paper.” When they occasionally engage with the readily available evidence, additional problems with BINs are immediately found. For example, for BIN “BOLD:AAG8211” the authors state under “other material” that “BIOUG07751-A11 (Malaise-trapped) from Honduras was examined and considered conspecific, and it is included as a paratype. There are dozens of specimens in the same BIN that are from widespread locations across Canada and the USA. Several of these were examined and they differ significantly in color from the Mesoamerican specimens and are not considered conspecific. See BOLD for details.” In particular, the North American specimens from Florida should have been consulted before delimiting the Costa Rican species, but these specimens are nevertheless neglected.

We suspect that Sharkey et al. (2021) overlooked the problems with delimiting species with COI because they relied too much on BINs supplied by BOLD Systems. As in all scientific studies, it is important that authors analyze the underlying data thoroughly in order to present robust conclusions. For species descriptions involving molecular data, the first step is to evaluate the decisiveness of the data by applying different species delimitation algorithms (Virgilio et al., 2010; Carstens et al., 2013; Kekkonen and Hebert, 2014; Yeo et al., 2020). It also means avoiding publication of results obtained with algorithms that are not publicly available (in this case: RESL) and operate on a mixture of public and private data.

Evidence for the lack of decisiveness can also been seen when studying the composition of BOLD BINs as more specimens are added to BOLD. This information is available when one traces the “BIN history” of specimens. Here is the history for three BINs described as species in Sharkey et al.’s (2021) minimalist revision (see suppl. materials for details):

- BOLD:AAH8697 (*Heterogamus donstonei*) has 54 members of which 18 are public and 14 are stable (have only been in one BIN). One unstable specimen (BCLDQ0377) was originally placed in a fly BIN (BOLD:AAG1770: Diptera: Muscidae: *Lispocephala varians*; 5 June 2010) before moving on to the current wasp BIN (19 June 2010). The three remaining unstable barcodes were all originally placed into the current BIN in September 2012, but subsequently shifted into two BINs without public members (in May 2013). All three then shifted back into the current BIN on 8 August 2015.
- BOLD:AAV3035 (*Pneumagathis erythrogastra*, new combination in Sharkey et al. (2021)) has 14 members of which 12 are barcode compliant. Only two barcodes are stable. The remaining 10 have similar histories. They were first placed in a stonefly BIN (BOLD:AAC5216: Plecoptera: Chloroperlidae: *Alloperla severa*; March 2010) before shifting to a BIN that is no longer available (January 2011). It is unclear when these specimens shifted to the current BIN because this is not revealed by the delta view tool in the BOLD Systems workbench. The two other barcodes were first placed in BIN(s) that are no longer available. One (specimen H1170) then shifted to the current BIN in September 2012 but later shifted to a BIN with a single private member in May 2013. This specimen returned to the current BIN in June 2018. The other specimen (specimen H7621) shifted from the original placement (BIN has ceased to exist) to the current BIN in June 2018.
- BOLD:ABY5286 (*Chelonus scottmilleri*). The history for the 15 public barcodes is detailed in Figure 3. It involves two BINs that are “no longer available” (BOLD:AAA2380 and BOLD:AAK1017) and two others that could not be accessed as they are “awaiting compliance with metadata requirements” (BOLD:AAD2009 and BOLD:ABX7466). Note that the two currently valid BINs (BOLD:AAA7014 and BOLD:ABX5499) contain species from different subfamilies of Braconidae (Microgastrinae: *Apanteles anamartinezae* and Cheloninae: *Chelonus scottshawi*, respectively).

**Figure 3.**
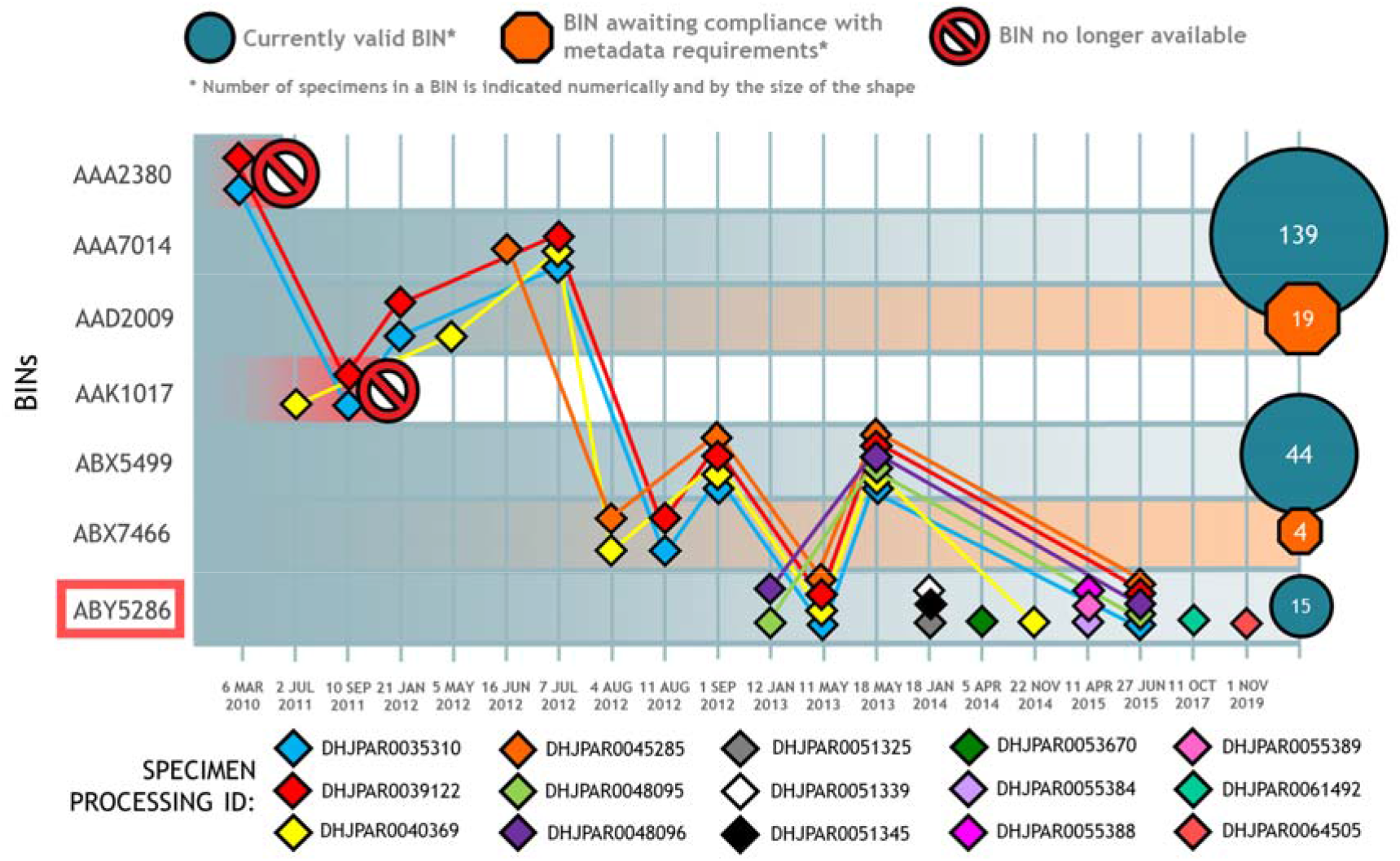
BIN membership through time for the 15 public barcodes in BOLD:ABY5286 (*Chelonus scottmilleri*). Two BINs are “no longer available”; two others cannot be accessed as they are “awaiting compliance with metadata requirements”; the two currently valid BINs contain species from different subfamilies (AAA7014 Microgastrinae: *Apanteles anamartinezae* and ABX5499 Cheloninae: *Chelonus scottshawi*). They contain 139 barcodes (all public) and 44 barcodes (although 45 listed as public), respectively.

All this highlights that relying heavily on BIN=Species for describing species is not advisable. Indeed, depending on the time of BIN description, two of Sharkey et al.’s (2021) wasp taxa would have been described as a fly or stonefly species, respectively. Similarly, biologists using BOLD as an identification tool should be aware that using BINs can lead to misleading conclusions. This makes it all the more important that BIN history can be traced easily. However, this is time-consuming in BOLD Systems. One can only reconstruct BIN evolution by tracing individual specimens – one at a time – and clicking through pages of changes over time; yet, this will still only reveal information on publicly available sequences. This is unfortunate because “taxon concepts” matter greatly in taxonomy (Franz, 2005; Meier, 2016; Packer et al., 2018) and it would be particularly straightforward to establish a “BIN-concept tracking” feature in BOLD Systems given that it is a relatively new database and only deals with one type of data.

The instability of species estimates obtained with barcodes is also evident when evaluating whether the original species diversity estimates in publications are consistent with species richness estimates based on the number of BINs in BOLD. We initially studied the data from publications that were highlighted in Sharkey et al. (2021). We then added the data for additional publications as long as they allowed for a direct comparison of species numbers (=authors provided a list of GenBank accession or BOLD sample IDs). We obtained the sequences in February-April 2021 using these numbers by querying GenBank, the Sequence-ID tool of GBIF, and BOLD Systems. Table 1 compares the number of species and BINs for those species that had molecular data. We find that, for example, the barcodes for the 10 species in *Astraptes* belong to 5 BINs (Hebert et al. 2004), the 32 species of *Belvosia* reported in Smith et al. (2006) belong to 20 BINs, and the barcodes in Smith et al. (2007) are in 61 BINs instead of the 73 species reported in the paper. These data illustrate how ambiguous COI can be when it comes to delimiting species. For the data from these studies, BINs are either not equal to species or the species reported in the publications were not species.

**Table 1:**
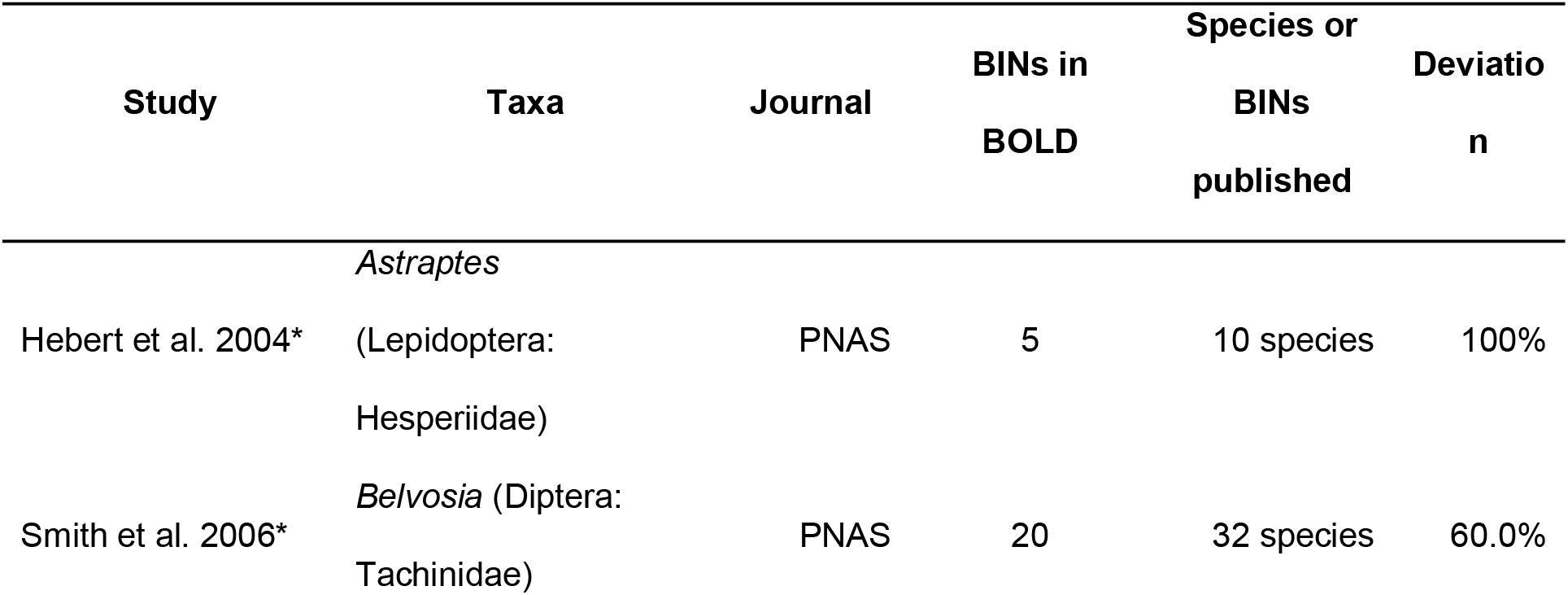
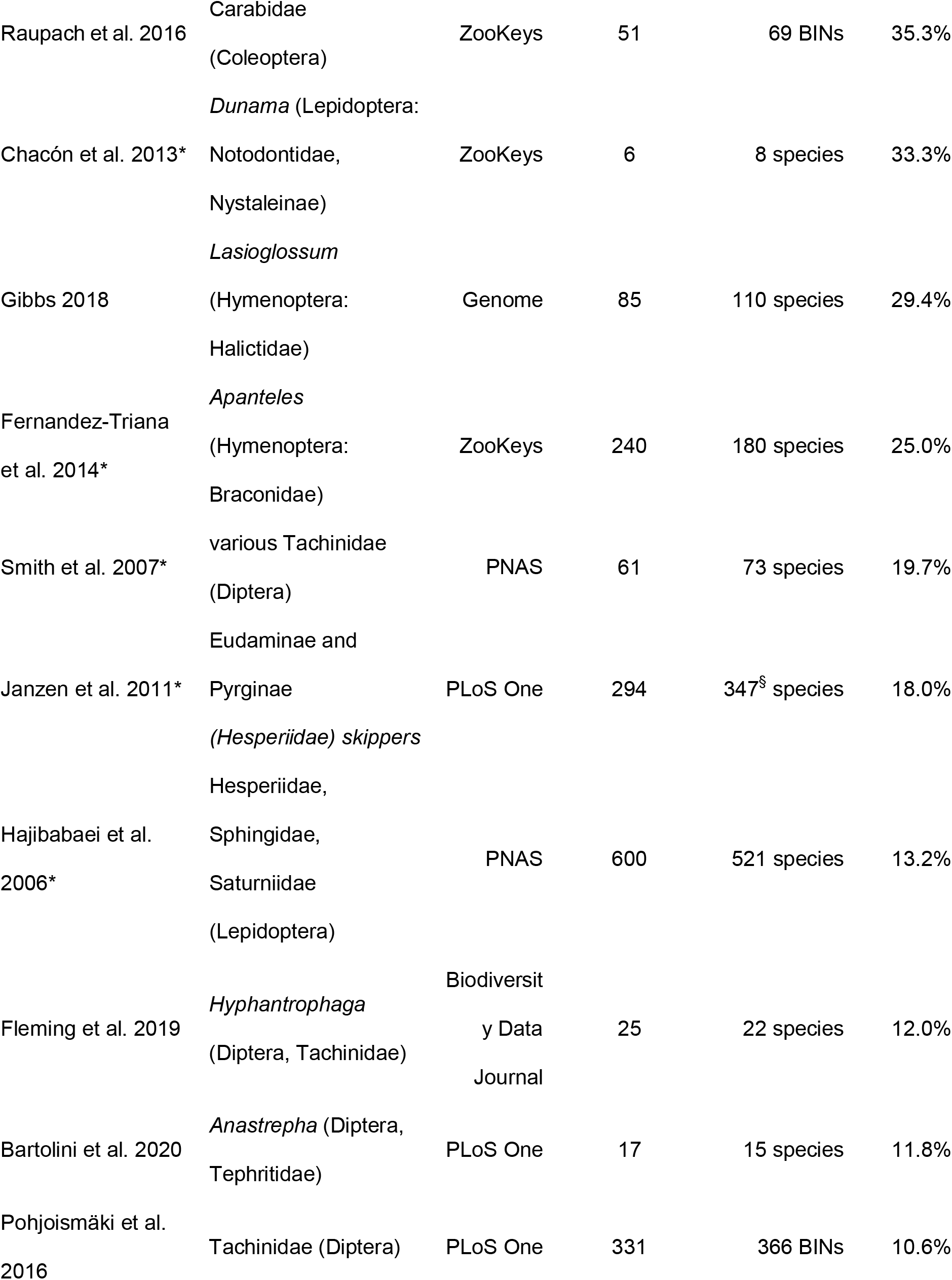
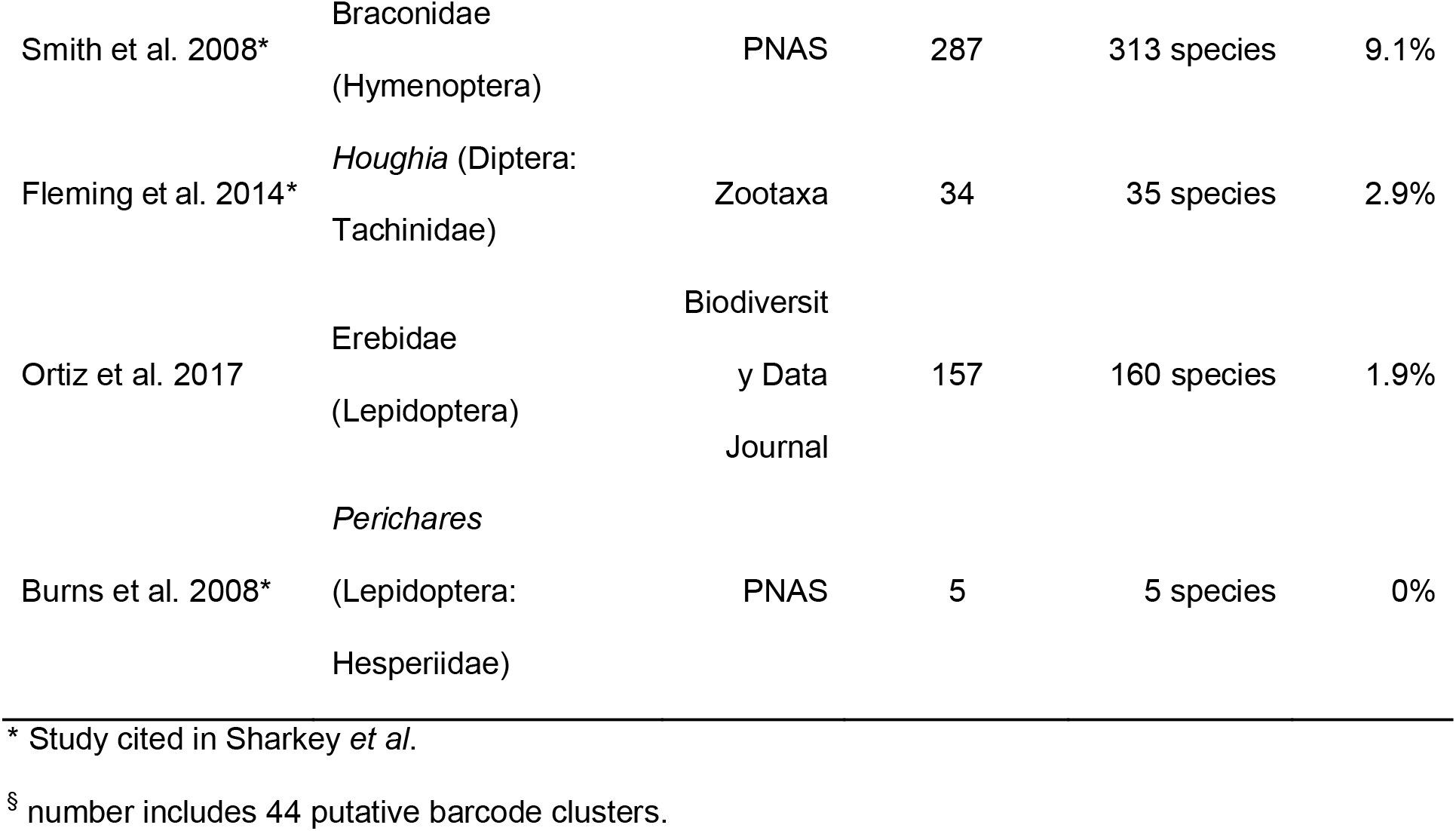
Comparison of published species and BIN numbers (April 2021).

In this context it is important to remember that there are good theoretical and empirical reasons why one should not expect *COI* clusters to be equal to species (Meier et al., 2006; Puillandre et al., 2012a; Puillandre et al., 2012b; Zhang et al., 2013). It has long been known that introgression, lineage sorting, and maternal inheritance of mt DNA can obfuscate the species-level signal in *COI* data (Will and Rubinoff, 2004; DeSalle et al., 2005). In addition, we know that for closely related taxa the COI protein is under strong stabilizing selection (Roe and Sperling, 2007; Kwong et al., 2012; Pentinsaari et al., 2016), which explains why most of the evolutionary change at the DNA level is synonymous and concentrated in 3^rd^ positions. Nucleotide fixation in these positions is likely caused by genetic drift; i.e., *COI* distances between BINs and closely related species mostly measure time of divergence which is only correlated but not identical to the probability of speciation.

This can be illustrated by the three datasets that we analyzed here. We find numerous BINs whose barcodes translate into identical amino acid sequences (Table S1: dataset 1: 11 cases involving 26 BINs; dataset 2: 21 cases and 49 BINs; dataset 3: 28 cases and 57 BINs – see supplementary materials for details) and the proportion in other taxa can be as high as 80-90% (Kwong et al., 2012). These BINs have no known, functionally significant biological differences because the biology of these wasp “species” is not at all affected by which triplet codons are used to code for identical proteins. Given that most biologists associate speciation with the origin of biologically meaningful differences, describing such BINs as species rests on the assumption that the correlation between time of divergence and the origin of new species is sufficiently strong that biologically meaningful differences will later be found. However, most biologists are rightfully skeptical of results based on correlations. Correlations warrant more exploration; i.e., barcode clusters should only be treated as unnamed first-pass groupings until they are validated with other data before being described as species.

An even more serious concern is the non-random nature of the error that is caused by faulty species delimitation with COI barcodes. Species originating quickly via an adaptive radiation will be systematically lumped while old species will be systematically split. Evidence for these problems can be found throughout BOLD. Anyone who has worked with the database will have found many cases where one BIN contains sequences that were identified as coming from multiple species. Conversely, it is common to find species that have barcodes in multiple BINs. This can be due to misidentification or genuine lumping/splitting. Fortunately, there are taxa for which there is enough morphological, genetic, and behavioral data that species boundaries are well understood. For these, one can test whether the sequences of recently diverged species end up in one BIN and the sequences for old species are found in several BINs. Such data are available, for example, for many species of Sepsidae (Diptera) (Puniamoorthy et al., 2009; Puniamoorthy et al., 2010; Tan et al., 2010; Ang et al., 2013a; Ang et al., 2013b; Araujo et al., 2014; Rohner et al., 2014; Ang et al., 2017). Known pairs of closely related species that are distinct with regard to morphology and behavior are routinely found lumped into the same BIN (e.g., *Sepsis neocynipsea* and *S. cynipsea*: BOLD:AAC2855; *Sepsis orthocnemis* and *S. fulgens*: BOLD:AAJ7599; *Themira lucida and T. flavicoxa*: BOLD:AAD7140). *Sepsis punctum* is an example for the opposite. The populations in North America and Europe can interbreed but are split into two BINs (BOLD:AAG5639; BOLD:ACS2531). If taxonomists had applied BIN=Species as a default for delimiting these species, several putatively young species would have been overlooked and some old species would have been split. Some will argue that this degree of error is acceptable, but we would counter that the non-random nature of the error is a major concern because it systematically misinforms about speciation.

## (2) Can species diagnoses consist of only a consensus barcode?

One of the key proposals of Sharkey et al. (2021) is that newly described species can be quickly and efficiently diagnosed using only one *COI* consensus barcode. Indeed, approximately 2/3rds of all newly described species in Sharkey et al. (2021) have such diagnoses. This approach to diagnosis would work if BINs=Species and there were no named species that were described based on morphological evidence. However, both conditions are not met and the minimalist revision thus struggles with implementing the “diagnosis=consensus barcode” concept. Indeed, one compromise follows another one until it becomes clear that morphological data remain indispensable for many species and highly desirable for others:

1. Several BINs contain more than one species. The consensus barcodes for 11 species thus have to be amended by brief morphological diagnoses (e.g. *Bracon tihisiaboshartae* can be differentiated from *B. alejandromasisi* by the color of the metasomal terga: entirely yellow in *B. tihisiaboshartae* and partly black in *B. alejandromasisi*”). The brevity of these amendments for all 11 species is reminiscent of species descriptions from the 18^th^ and 19^th^ century, and it appears very unlikely that these diagnoses will stand the test of time for species in a genus like *Bracon*, which “is an enormous, polyphyletic, cosmopolitan genus with thousands of undescribed species (Sharkey et al., 2021).” Such superficial diagnoses have the potential to become a major “superficial description impediment” in future.
2. Some species within the revision lack molecular data. Three new species are described based on short morphological diagnoses: “Differing from the five other known species (two undescribed and not included here) in many characters including a hypopygium that is much more convex than the others. Body length 7.1 mm. Other details can be gleaned from the images in Fig. 367”. The description of these species based on morphology means that all new, congeneric species delimited with DNA barcodes also need morphological information in the diagnoses. Otherwise, it cannot be shown that the new species are different. Sharkey et al. (2021) again amend the consensus barcodes with very short morphological circumscriptions, which are unlikely to be sufficiently informative once more Central American species are studied.
3. Two new species are described based on *COI* sequences that are too short to be barcodes according to the thresholds set by BOLD. Therefore neither has its own BIN identifier. Sharkey et al. (2021) proceed to describe one species based on a consensus mini-barcode which is again amended with morphological information ("Unique within Rogadinae, this species possesses a strongly petiolate (clavate) hind femur. The patchily patterned wings are also diagnostic within the *Colastomion-Cystomastax* group.”). For the other, the sequence information is ignored and only morphological information is provided: “The COI barcode has only 293 bases and therefore was not given a BIN. This species can be distinguished from all other species of *Zacremnops* by red coloration restricted to the metapleuron (i.e., not on the propodeum) in combination with the melanic hind tarsus.”
4. A special challenge to the “diagnosis=consensus barcode” concept are the 22 BINs that contain non-Costa Rican specimens. These BINs and their consensus barcodes were computed based on all data, but Sharkey et al.’s (2021) revision only covers the Costa Rican species. This is problematic because the diagnoses based on consensus barcodes are only correct when all specimens from outside of Costa Rica belong to the Costa Rican species. This is unlikely for two reasons. Firstly, some of the non-Costa Rican specimens are from geographically distant localities (e.g., Oklahoma, Kentucky, Mexico, Columbia, Bolivia, Argentina). Secondly, Sharkey et al. (2021) occasionally conclude based on images for non-Costa Rican specimens that some BINs contain >2 species. Sometimes, this leads to a diagnosis that consists of a consensus barcode and a geographic amendment (e.g., *Phanerotoma angelsolisi* consists of those specimens in BOLD:AAG8211 that are “Restricted to Central America”). In other cases, the amendment is morphological (*Bracon mariamartachavarriae* consists of those specimens of BOLD:AAV6295 that have a “Terminal flagellomeres black, concolorous with basal flagellomeres”). However, the validity of the diagnoses is in question because it is unclear whether critical specimens were studied (e.g., specimens from Florida for BOLD:AAG8211; specimen from Argentina for BOLD:AAV6295). It is also unclear why Sharkey et al. (2021) use consensus barcode for entire BINs when the authors only want the species to consist of the Costa Rican specimens in a BIN. For BINs with wide distributions, such diagnoses are very unlikely to pertain to the Costa Rican species. Note that there are other cases where the diagnoses consisting only of a consensus barcode are almost certainly incorrect because the authors indicate that the BINs appear to contain more than one species (e.g., BOLD:AAH8705 and BOLD:ABZ7672).

There are additional oddities with regard to the use of consensus barcodes as diagnoses, which highlight that relying only on one type of data is ill-advised. BOLD:ABX6701 was described by Sharkey et al. (2021) as *Plesiocoelus vanachterbergi*. The consensus barcode includes two single indels and was presumably obtained from the seven “barcode-compliant” sequences in BOLD Systems (as of April 18, 2021). Only one of the seven barcodes is translatable to amino acids with the remaining six having deletions. Such sequences are typical for pseudogenes, and it is conceivable that *P. vanachterbergi* was delimited and is now diagnosed based on paralogs. In addition, the consensus barcode for *Pseudorhysipolis mailyngonzalezae* contains 104 indels, which may be related to the fact that the BOLD fasta download for this BIN consists of data for multiple genes (*COI*, *28S*, *16S*, *Ef1a*).

We here discuss these issues because diagnoses are critical elements of species descriptions. They render species hypotheses testable (Ahrens et al., 2021). A diagnosis consisting of only a consensus barcode cannot be tested with morphological data and vice versa. This is one reason why integrative taxonomy is preferred over other approaches. Sharkey et al.’s (2021) reliance on consensus barcodes is particularly problematic because they are static summaries of dynamically evolving barcode databases. More sequencing will almost certainly reveal synonymous nucleotide changes in barcodes for the more than 200 species of braconids for which Sharkey et al. (2021) only provide consensus barcodes as diagnoses; i.e., these new polymorphisms will falsify the diagnoses. By omitting morphological information for these species, Sharkey et al. also created a parallel taxonomy that does not allow for cross-referencing the new species to the described fauna (Ahrens et al., 2021; Zamani et al., 2021). It furthermore interferes with the work of those biologists, policymakers, and citizen scientists in biodiverse countries who have no access to molecular data (Zamani et al., 2021).

Our discussion of consensus barcodes as diagnoses also highlights Sharkey et al.’s (2021) confusing relationship to morphological evidence. On the one hand, Sharkey et al. (2021) argue that morphological evidence should not be used for braconid taxonomy. On the other hand, a large number of diagnoses contain morphological information, and in many cases the morphological characters are essential for the validity of the species. Morphology was presumably also used to identify those BINs that consist of multiple species. Despite this, no integrative process is described and the minimalist revision failed to provide proper morphological diagnoses/description for many species.

## (3) Should BINs obtained from BOLD Systems be used as species estimates?

There are three reasons why the BINs in the “minimalist revision” do not meet the minimum standards for reproducible science. All three are related to open science issues within BOLD Systems. Firstly, the algorithm for calculating BINs (RESL: Refined Single Linkage) is not publicly available. Secondly, it is unclear how the underlying sequences were aligned. Thirdly, BINs are calculated based on public and private barcodes:

### RESL: Refined Single Linkage

The code for this algorithm that computes BINs is proprietary. This makes it impossible to vary parameters such as the pairwise distances used for the initial clustering, the thresholds used for merging neighbors, and the inflation parameters for cluster refinement. This lack of user control can give the impression that BIN boundaries are well defined, but this is achieved by not allowing users to vary parameters; i.e., BINs are more akin to means that lack standard deviations, or a tree with nodes that lack information on support. Barcode data should always be analyzed using several species delimitation algorithms in order to investigate how robust different barcode clusters are.

Overall, we know very little about RESL’s performance because it was simultaneously proposed and implemented based on only eight small datasets that comprised 18,843 barcodes (Ratnasingham and Hebert, 2013). The optimal clustering thresholds for these datasets varied from 0.7% to 1.8% and eventually “2.2% was adopted as it represents the upper 99% confidence limit for the optimal thresholds in the eight test datasets (x□=1.26), SD =0.40)” (Ratnasingham and Hebert, 2013). The same eight datasets were also analyzed with four other species delimitation algorithms (ABGD, CROP, GMYC, jMOTU). Overall, RESL and GMYC performed best with 89% congruence with morphology. Note, however, that this level of congruence was only achieved after RESL was applied to the same eight datasets for which it was optimized.

All remaining comparisons of RESL to other species delimitation algorithms are compromised because RESL can only be used within BOLD Systems and the other delimitation algorithms cannot be applied to BOLD System’s private data. This means that the comparative performance of RESL remains unclear. However, it is unlikely that it is a superior method. As discussed by Ratnasingham and Hebert (2011), the algorithm was developed with speed and scalability in mind and the authors anticipated that “[f]uture research will undoubtedly reveal analytical approaches that are better at recognizing species boundaries from sequence information than RESL.” This means that taxonomic revisions should not overly rely on BINs. Instead, the available data should be analyzed using a range of species delimitation algorithms. As shown earlier, this would have provided important insights into the robustness of BINs. Note that the use of different analysis tools also allows users to gain control over the quality of the alignments. This is important because this is another open science issue in Bold Systems given that the “parameters for BOLD’s alignment of sequences are not publicly available” (Nugent et al., 2020) and cannot be changed.

### Private data

The third reason why Sharkey et al. should not have relied on BINs is that they are calculated based on public and private barcodes. The ratio of public and private barcodes is difficult to determine because the relevant numbers vary between what is listed on BOLD Systems under “Identification engine” and “Taxonomy”, and what can be downloaded from BOLD Systems via the “Public Data Portal”. We here estimate that approximately one quarter of all data are private. In order to obtain this estimate, we downloaded the data for each taxonomic category listed under Animals in BOLD Systems’ “Taxonomy”. Given the large size of the Arthropoda and Insecta datasets, sub-taxa were downloaded separately. Downloading was done either directly from the web browser or by using BOLD Systems’ API (27 March-11 April 2021). Overall, we obtained 6,165,928 COI barcodes >500 bp (6,508,300 overall) of the 8,480,276 (18 April 2021) barcodes that are listed as “All Barcode Records on BOLD”. We estimate that obtaining these data and setting up the facilities and bioinformatics pipelines cost >200 million C$, given that one project alone (Barcode500K: 2010-2015) had a budget of 125 million C$ (https://ibol.org/programs/barcode-500k/). It is very likely that some of these funds were also used for generating private data. This includes private BINs that have remained private for many years.

### Use of potentially misleading tools

The value of Sharkey et al.’s (2021) minimalist revision is not only diminished because it relies too heavily on BINs, but the authors were also undiscerning in their choice of tools. Indeed, many are considered “deadly sins of DNA barcoding” in Collins and Cruickshank (2013). This included the use of Neighbor-Joining (NJ) trees which can yield biased results when the input order of taxa is not randomized (Takezaki, 1998). BOLD Systems’ K2P NJ trees feature prominently in Sharkey et al. although they lack support values, information on how tie-trees are broken, and use with K2P an inappropriate evolutionary model for *COI* evolution (Collins et al., 2012; Srivathsan and Meier, 2012). Note that this model is not even used by RESL when calculating BINs. Many authors – including Sharkey et al. (2021) – argue that the use of K2P NJ trees is acceptable because they are “only” used for visualization. However, poorly designed visualization tools can misinform about the validity and support for groupings.

All of the issues herein should have been addressed during manuscript review. This is why we urge reviewers, journals, and grantors to be more rigorous when reviewing barcoding studies and projects. A lenient attitude toward BIN data and BOLD has resulted in scientific publications that do not meet the minimum standards for reproducible science. Some journals seem to neglect their own data and code accessibility standards or else papers that are critically dependent on BINs would not be published. Similarly, a major funder of DNA barcoding is Genome Canada, which has an explicit data sharing policy that contradicts how RESL and alignment parameters are handled (see https://www.genomecanada.ca/en/programs/guidelines-and-policies).

## Final Remarks and Suggestions

Sharkey et al. (2021) are pessimistic about new 21^st^ century solutions to the dark-taxon impediment. We do not share this sentiment. Large throughput imaging and sequencing are already available, these are not vague promises for the future (Hebert et al., 2018; Ärje et al., 2020; Srivathsan et al., 2021; Wührl et al., 2021). Similarly, data can now be analyzed with increasingly sophisticated algorithms that will provide taxonomists with a solid foundation for species descriptions that can be based on multiple sources of data (Puillandre et al., 2012b; Hartop et al., 2021). These data will be particularly suitable for generating automatic species descriptions that are resilient and future ready. Therefore, this is the worst time to propose “minimalist revisions” that trade scientific rigor for speed. We would like to conclude our critique of Sharkey et al. (2021) with some suggestions for BOLD:

- The RESL algorithm must be published and access must be provided to the private data, or these data must be excluded from BIN calculation.
- BOLD Systems should implement additional clustering algorithms for barcode data. This would provide users with much needed information on which barcode clusters are congruent across methods.
- BOLD Systems should include a BIN tracking tool for “BIN concepts.” It is essential for validating the results of studies that used BINs computed in the past.
- The relationship between BINs and species in BOLD Systems should be revisited. The original BOLD Systems publication highlighted the workbench character of the database. BINs known to contain several verified species were supposed to be labeled with decimal numbers (“BOLD:AAB2314.1”). However, this option has been largely ignored. This means that the BIN matches of searches within BOLD Systems or GBIF’s SequenceID lead to BINs, even if they are known to contradict species boundaries. BINs containing sequences for multiple species should be flagged.

## Acknowledgements

We would like to thank Drs Roderic D. M. Page, Steve Marshall, and one additional colleague for providing valuable comments on the manuscript. This work was supported by a Ministry of Education grant on biodiversity discovery (R-154-000-A22-112).

## Data accessibility statement

All alignments and scripts are available from https://figshare.com/articles/dataset/BOLD_braconid_datasets/14452992.

## References

Ahrens, D., Ahyong, S. T., Ballerio, A., Barclay, M. V. L., Eberle, J., Espelund, M., Huber, B. A., Mengual, X., Pacheco, T. L., Peters, R. S., Rulik, B., Vaz-de-Mello, F., Wesener, T., K, rell, F. T. 2021. Is it time to describe new species without diagnoses? - A comment on Sharkey et al. (2021). Zenodo, 10.5281/zenodo.4899151.

Ang, Y., Puniamoorthy, J., Pont, A. C., Bartak, M., Blanckenhorn, W. U., Eberhard, W. G., Puniamoorthy, N., Silva, V. C., Munari, L., Meier, R. 2013a. A plea for digital reference collections and other science-based digitization initiatives in taxonomy: Sepsidnet as exemplar. Systematic Entomology 38, 637–644.

Ang, Y., Rajaratnam, G., Su, K. F., Meier, R. 2017. Hidden in the urban parks of New York City: *Themira lohmanus*, a new species of Sepsidae described based on morphology, DNA sequences, mating behavior, and reproductive isolation (Sepsidae, Diptera). ZooKeys, 95–111.

Ang, Y. C., Wong, L. J., Meier, R. 2013b. Using seemingly unnecessary illustrations to improve the diagnostic usefulness of descriptions in taxonomy-a case study on *Perochaeta orientalis* (Diptera, Sepsidae). Zookeys, 9–27.

Araujo, D., Tuan, M., Yew, J., Meier, R. 2014. Analysing small insect glands with UV-LDI MS: high-resolution spatial analysis reveals the chemical composition and use of the osmeterium secretion in *Themira superba* (Sepsidae: Diptera). Journal of Evolutionary Biology 27, 1744–1750.

Ärje, J., Melvad, C., Jeppesen, M. R., Madsen, S. A., Raitoharju, J., Rasmussen, M. S., Iosifidis, A., Tirronen, V., Gabbouj, M., Meissner, K., Høye, T. T. 2020. Automatic image-based identification and biomass estimation of invertebrates. 11, 922–931.

Bartolini, I., Rivera, J., Nolazco, N., Olortegui, A. 2020. Towards the implementation of a DNA barcode library for the identification of Peruvian species of *Anastrepha* (Diptera: Tephritidae). Plos One 15.

Bergsten, J., Bilton, D. T., Fujisawa, T., Elliott, M., Monaghan, M. T., Balke, M., Hendrich, L., Geijer, J., Herrmann, J., Foster, G. N., Ribera, I., Nilsson, A. N., Barraclough, T. G., Vogler, A. P. 2012. The effect of geographical scale of sampling on DNA barcoding. Systematic Biology 61, 851–869.

Burns, J. M., Janzen, D. H., Hajibabaei, M., Hallwachs, W., Hebert, P. D. N. 2008. DNA barcodes and cryptic species of skipper butterflies in the genus *Perichares* in Area de Conservacion Guanacaste, Costa Rica. Proceedings of the National Academy of Sciences of the United States of America 105, 6350–6355.

Carstens, B. C., Pelletier, T. A., Reid, N. M., Satler, J. D. 2013. How to fail at species delimitation. Molecular Ecology 22, 4369–4383.

Chacon, I. A., Janzen, D. H., Hallwachs, W., Sullivan, J. B., Hajibabaei, M. 2013. Cryptic species within cryptic moths: new species of *Dunama* Schaus (Notodontidae, Nystaleinae) in Costa Rica. Zookeys, 11–45.

Collins, R., Cruickshank, R. 2013. The seven deadly sins of DNA barcoding. Molecular Ecology Resources 13, 969–975.

Collins, R. A., Boykin, L. M., Cruickshank, R. H., Armstrong, K. F. 2012. Barcoding’s next top model: an evaluation of nucleotide substitution models for specimen identification. Methods in Ecology and Evolution 3, 457–465.

DeSalle, R., Egan, M. G., Siddall, M. 2005. The unholy trinity: taxonomy, species delimitation and DNA barcoding. Philosophical Transactions of the Royal Society B-Biological Sciences 360, 1905–1916.

Fernandez-Triana, J. L., Whitfield, J. B., Rodriguez, J. J., Smith, M. A., Janzen, D. H., Hallwachs, W. D., Hajibabaei, M., Burns, J. M., Solis, M. A., Brown, J., Cardinal, S., Goulet, H., Hebert, P. D. N. 2014. Review of *Apanteles* sensu stricto (Hymenoptera, Braconidae, Microgastrinae) from Area de Conservacion Guanacaste, northwestern Costa Rica, with keys to all described species from Mesoamerica. Zookeys, 1–565.

Fleming, A. J., Wood, D. M., Smith, M. A., Dapkeyl, T., Hallwachs, W., Janzen, D. 2019. Twenty-two new species in the genus *Hyphantrophaga* Townsend (Diptera: Tachinidae) from Area de Conservacion Guanacaste, with a key to the species of Mesoamerica. Biodiversity Data Journal 7.

Fleming, A. J., Wood, D. M., Smith, M. A., Hallwachs, W., Janzen, D. H. 2014. Revision of the New World species of *Houghia* Coquillett (Diptera, Tachinidae) reared from caterpillars in Area de Conservacion Guanacaste, Costa Rica. Zootaxa 3858, 1–90.

Forum, W. E. 2020. World Economic Forum. The Global Risks Report 2020.

Franz, N. M. 2005. On the lack of good scientific reasons for the growing phylogeny/classification gap. Cladistics 21, 495–500.

Gibbs, J. 2018. DNA barcoding a nightmare taxon: assessing barcode index numbers and barcode gaps for sweat bees. Genome 61, 21–31.

Hajibabaei, M., Janzen, D. H., Burns, J. M., Hallwachs, W., Hebert, P. D. N. 2006. DNA barcodes distinguish species of tropical Lepidoptera. Proceedings of the National Academy of Sciences of the United States of America 103, 968–971.

Hartop, E., Srivathsan, A., Ronquist, F., Meier, R. 2021. Large-scale Integrative Taxonomy (LIT): resolving the data conundrum for dark taxa. bioRxiv 10.1101/2021.1104.1113.439467.

Hausmann, A., Krogmann, L., Peters, R. S., Rduch, V., Schmidt, S. 2020. GBOL III: Dark taxa. Barcode Bulletin 10.

Hebert, P. D., Braukmann, T. W., Prosser, S. W., Ratnasingham, S., DeWaard, J. R., Ivanova, N. V., Janzen, D. H., Hallwachs, W., Naik, S., Sones, J. E. 2018. A Sequel to Sanger: amplicon sequencing that scales. 19, 1–14.

Hebert, P. D. N., Penton, E. H., Burns, J. M., Janzen, D. H., Hallwachs, W. 2004. Ten species in one: DNA barcoding reveals cryptic species in the Neotropical skipper butterfly *Astraptes fulgerator*. Proceedings of the National Academy of Sciences of the United States of America 101, 14812–14817.

Janzen, D. H., Hallwachs, W., Burns, J. M., Hajibabaei, M., Bertrand, C., Hebert, P. D. N. 2011. Reading the complex skipper butterfly fauna of one tropical place. Plos One 6.

Kekkonen, M., Hebert, P. D. N. 2014. DNA barcode-based delineation of putative species: efficient start for taxonomic workflows. Molecular Ecology Resources 14, 706–715.

Kwong, S., Srivathsan, A., Vaidya, G., Meier, R. 2012. Is the COI barcoding gene involved in speciation through intergenomic conflict? Molecular Phylogenetics and Evolution 62, 1009–1012.

May, R. M. 2011. Why worry about how many species and their loss? PLoS Biology 9.

Meier, R. 2016. Citation of taxonomic publications: the why, when, what and what not. Systematic Entomology 42, 301–304.

Meier, R., Shiyang, K., Vaidya, G., Ng, P. K. L. 2006. DNA barcoding and taxonomy in diptera: A tale of high intraspecific variability and low identification success. Systematic Biology 55, 715–728.

Nugent, C. M., Elliott, T. A., Ratnasingham, S., Adamowicz, S. J. 2020. Coil: an R package for cytochrome c oxidase I (COI) DNA barcode data cleaning, translation, and error evaluation. Genome 63, 291–305.

Ortiz, A. S., Rubio, R. M., Guerrero, J. J., Garre, M. J., Serrano, J., Hebert, P. D. N., Hausmann, A. 2017. Close congruence between Barcode Index Numbers (bins) and species boundaries in the Erebidae (Lepidoptera: Noctuoidea) of the Iberian Peninsula. Biodiversity Data Journal 5.

Packer, L., Monckton, S. K., Onuferko, T. M., Ferrari, R. R. 2018. Validating taxonomic identifications in entomological research. Insect Conservation and Diversity 11, 1–12.

Pentinsaari, M., Salmela, H., Mutanen, M., Roslin, T. 2016. Molecular evolution of a widely-adopted taxonomic marker (COI) across the animal tree of life. Scientific Reports 6, 35275.

Pohjoismaki, J. L. O., Kahanpaa, J., Mutanen, M. 2016. DNA barcodes for the Northern European tachinid flies (Diptera: Tachinidae). Plos One 11.

Puillandre, N., Brouillet, S., Achaz, G. 2021. ASAP: assemble species by automatic partitioning. Molecular Ecology Resources 21, 609–620.

Puillandre, N., Lambert, A., Brouillet, S., Achaz, G. 2012a. ABGD, Automatic Barcode Gap Discovery for primary species delimitation. Molecular Ecology 21, 1864–1877.

Puillandre, N., Modica, M. V., Zhang, Y., Sirovich, L., Boisselier, M.-c., Cruaud, C., Holford, M., Samadi, S. 2012b. Large-scale species delimitation method for hyperdiverse groups. Molecular Ecology 21, 2671–2691.

Puniamoorthy, N., Ismail, M. R. B., Tan, D. S. H., Meier, R. 2009. From kissing to belly stridulation: comparative analysis reveals surprising diversity, rapid evolution, and much homoplasy in the mating behaviour of 27 species of sepsid flies (Diptera: Sepsidae). Journal of Evolutionary Biology 22, 2146–2156.

Puniamoorthy, N., Kotrba, M., Meier, R. 2010. Unlocking the “Black box”: internal female genitalia in Sepsidae (Diptera) evolve fast and are species-specific. BMC Evolutionary Biology 10.

Ratnasingham, S., Hebert, P. D. N. 2011. BOLD’s role in barcode data management and analysis: a response. Molecular Ecology Resources 11, 941–942.

Ratnasingham, S., Hebert, P. D. N. 2013. A DNA-based registry for all animal species: The Barcode Index Number (BIN) system. Plos One 8.

Raupach, M. J., Hannig, K., Moriniere, J., Hendrich, L. 2016. A DNA barcode library for ground beetles (Insecta, Coleoptera, Carabidae) of Germany: The genus *Bembidion* Latreille, 1802 and allied taxa. Zookeys, 121–141.

Roe, A. D., Sperling, F. A. H. 2007. Patterns of evolution of mitochondrial cytochrome c oxidase I and II DNA and implications for DNA barcoding. Molecular Phylogenetics and Evolution 44, 325–345.

Rohner, P. T., Ang, Y., Lei, Z., Puniamoorthy, N., Blanckenhorn, W. U., Meier, R. 2014. Genetic data confirm the species status of *Sepsis nigripes* Meigen (Diptera: Sepsidae) and adds one species to the Alpine fauna while questioning the synonymy of *Sepsis helvetica* Munari. Invertebrate Systematics 28, 555–563.

Smith, M. A., Rodriguez, J. J., Whitfield, J. B., Deans, A. R., Janzen, D. H., Hallwachs, W., Hebert, P. D. N. 2008. Extreme diversity of tropical parasitoid wasps exposed by iterative integration of natural history, DNA barcoding, morphology, and collections. Proceedings of the National Academy of Sciences of the United States of America 105, 12359–12364.

Smith, M. A., Wood, D. M., Janzen, D. H., Hallwachs, W., Hebert, P. D. N. 2007. DNA barcodes affirm that 16 species of apparently generalist tropical parasitoid flies (Diptera, Tachinidae) are not all generalists. Proceedings of the National Academy of Sciences of the United States of America 104, 4967–4972.

Smith, M. A., Woodley, N. E., Janzen, D. H., Hallwachs, W., Hebert, P. D. N. 2006. DNA barcodes reveal cryptic host-specificity within the presumed polyphagous members of a genus of parasitoid flies (Diptera: Tachinidae). Proceedings of the National Academy of Sciences of the United States of America 103, 3657–3662.

Srivathsan, A., Hartop, E., Puniamoorthy, J., Lee, W. T., Kutty, S. N., Kurina, O., Meier, R. 2019. Rapid, large-scale species discovery in hyperdiverse taxa using 1D MinION sequencing. BMC Biology 17, 96.

Srivathsan, A., Lee, L., Katoh, K., Hartop, E., Kutty, S. N., Wong, J., Yeo, D., Meier, R. J. b. 2021. MinION barcodes: biodiversity discovery and identification by everyone, for everyone. BioRxiv.

Srivathsan, A., Meier, R. 2012. On the inappropriate use of Kimura-2-parameter (K2P) divergences in the DNA-barcoding literature. Cladistics 28, 190–194.

Stamatakis, A. J. B. 2014. RAxML version 8: a tool for phylogenetic analysis and post-analysis of large phylogenies. 30, 1312–1313.

Takezaki, N. 1998. Tie trees generated by distance methods of phylogenetic reconstruction. Molecular Biology and Evolution 15, 727–737.

Tan, D. S., Ang, Y., Lim, G. S., Ismail, M. R. B., Meier, R. 2010. From ‘cryptic species’ to integrative taxonomy: an iterative process involving DNA sequences, morphology, and behaviour leads to the resurrection of Sepsis pyrrhosoma (Sepsidae: Diptera). Zoologica Scripta 39, 51–61.

Virgilio, M., Backeljau, T., Nevado, B., De Meyer, M. J. B. b. 2010. Comparative performances of DNA barcoding across insect orders. BMC Bioinformatics 11, 1–10.

Will, K. W., Rubinoff, D. 2004. Myth of the molecule: DNA barcodes for species cannot replace morphology for identification and classification. Cladistics 20, 47–55.

Wührl, L., Pylatiuk, C., Giersch, M., Lapp, F., von Rintelen, T., Balke, M., Schmidt, S., Cerretti, P., Meier, R. 2021. DiversityScanner: Robotic discovery of small invertebrates with machine learning methods. 2021.2005.2017.444523.

Yeo, D., Srivathsan, A., Meier, R. 2020. Longer is not always better: Optimizing barcode length for large-scale species discovery and Identification. Systematic Biology 69, 999–1015.

Zamani, A., Vahtera, V., Sääksjärvi, I. E., Scherz, M. D. 2021. The omission of critical data in the pursuit of ‘revolutionary’ methods to accelerate the description of species. Systematic Entomology 46, 1–4.

Zhang, J., Kapli, P., Pavlidis, P., Stamatakis, A. 2013. A general species delimitation method with applications to phylogenetic placements. Bioinformatics 29, 2869–2876.

