## Supplementary methods for "A re-analysis of the data in Sharkey et al.’s (2021) minimalist revision reveals that BINs do not deserve names, but BOLD Systems needs a stronger commitment to open science" (from another version of this preprint: version 3, 2021-05-23)

**Supplementary Information**

**Materials & Methods**

*Data retrieval*. Barcode data were extracted from BOLD for the three datasets via the following approaches BIN numbers were added to the sequence headers of each barcode for all datasets:

- Dataset 1: New BINs from Sharkey et al. (2021): A list of BINs were compiled from the Sharkey et al. (2021) paper. The BINs were then retrieved from BOLD via the following url: http://www.BOLD Systems.org/index.php/API_Public/sequence?bin=BOLD:**<BIN>**
- Dataset 2: All BINs from NJ Trees: A list of BINs were compiled from the NJ trees in the supplementary of Sharkey et al. (2021). The BINs were then retrieved from BOLD via the following url: http://www.BOLD Systems.org/index.php/API_Public/sequence?bin=BOLD:**<BIN>**
- Dataset 3: All subfamily BINs from BOLD: The subfamily names included in the paper were used as search terms in BOLD System’s public data portal. The barcodes were downloaded in FASTA format and compiled.

*Species delimitation*. The sequences in all three datasets were first aligned with MAFFT v7 (Katoh and Standley 2013) using a gap opening penalty of 5.0. Dataset 3 initially produced an untranslatable alignment with multiple indels. Sequences with a large number of internal gaps were separated from the other barcodes, with the remaining processed using a gap opening penalty of 5.0, while the --add function was used to add the problematic barcodes to the alignment using the default gap opening penalty. All alignments and scripts are available from https://figshare.com/articles/dataset/BOLD_braconid_datasets/14452992. The aligned barcodes were then processed using three species delimitation algorithms that represent both distance and tree-based approaches. Objective clustering under TaxonDNA (Meier et al. 2006) was performed using a range of distance thresholds (1-5% p-distance) common in the *COI* barcoding literature for Metazoa (cluster algorithm in TaxonDNA: Meier et al. 2006, 2016, Ratnasingham & Hebert 2013). ASAP (Puillandre et al. 2021) was performed using simple distances (d -3) and the top five partitions (lowest p-values) were used in this study. A maximum likelihood tree was generated with RAxML v.8 (Stamatakis 2014) via its rapid hill-climbing algorithm (-f d) under the GTRCAT model on CIPRES (Miller et al. 2011). The resulting best tree was processed with single-rate PTP (--single) (Zhang et al. 2013) using the mPTP scripts (Kapli et al. 2017).

Python scripts were used to identify which clusters from the above species delimitation algorithms and parameters were congruent with the BINs in a pairwise manner. The most congruent parameters for each delimitation algorithm (PTP, ASAP, objective clustering under TaxonDNA) were identified and cluster congruence visualized with nVenn diagrams (Pérez-Silva et al. 2018).

*Translation and amino acid distances*. The barcodes for the three datasets were aligned with TranslatorX (Abascal et al., 2010) using the invertebrate mitochondrial genetic code with the MAFFT method. The aligned amino acid sequences were processed using the UCLUST algorithm (Edgar 2010) with the cluster_fast command at -id 1.0 to obtain clusters of identical amino acid sequences. A Python script was then used to identify and quantify which of these identical clusters contained multiple BINs.

Table S1. List of BINs with identical amino acid sequences

*Dataset 1:*

| Case | BINs | Case | BINs |
| --- | --- | --- | --- |
| 1 | AAD7334, ABU7454 | 7 | AAA5374, AAB8975, ACK7801 |
| 2 | AAK1016, ABX5499, ABY5286, ACJ3551 | 8 | AAB1652, ABA9319, ACX5348 |
| 3 | AAB1658, ADB1219 | 9 | AAE0329, ADJ3083 |
| 4 | AAA5372, AAJ4092 | 10 | AAJ3968, ABZ7672 |
| 5 | ACJ4722, ACL5572 | 11 | ABZ9478, ACJ4626 |
| 6 | ADF4538, ADA8821 |  |  |

*Dataset 2:*

| Case | BINs | Case | BINs |
| --- | --- | --- | --- |
| 1 | AAB8975, ACK7801 | 12 | ACL5572, ACJ4722 |
| 2 | AAM5951, ACG8400, ACJ2495, ACO7975 | 13 | ADF4538, ADA8821 |
| 3 | AAK1016, ABX5499, ABY5286, ACJ3551 | 14 | ACJ5020, ADC9608 |
| 4 | ACN9714, ADC9450 | 15 | AAA5374, AAB8975, ACK7801, AEB4766 |
| 5 | AAB1658, ADB1219 | 16 | AAB1652, ABA9319, ACX5348 |
| 6 | AAD7334, ABU7454 | 17 | AAE0329, ADJ3083 |
| 7 | ACA4726, ACA4727, ADC9733, AEB4152 | 18 | AAJ3968, ABZ7672 |
| 8 | ACB1292, ACY8111 | 19 | ABZ9478, ACJ4626 |
| 9 | AAA5372, AAJ4092 | 20 | ACJ5330, ADC9429, ADC9430 |
| 10 | ABA9319, ABY3599 | 21 | ACK7467, AEF5717 |
| 11 | ACJ2111, ACN0949 |  |  |

*Dataset 3:*

| Case | BINs | Case | BINs |
| --- | --- | --- | --- |
| 1 | AAU8750, ACF4955, ADJ2714 | 15 | AAH8628, AAV7498 |
| 2 | AAG1434, ADA8867 | 16 | ACF4946, AAH8726 |
| 3 | AAD7425, AAU9842 | 17 | ACE1136, AAK5490 |
| 4 | AAH0622, ACQ8886 | 18 | ACD2708, AAJ1227 |
| 5 | ACE4571, AEA6054 | 19 | ACI8848, AAV7450 |
| 6 | AAG1395, ADE1516 | 20 | ACT1567, ACV2385 |
| 7 | ABY5568, AAD9345 | 21 | AEG0007, ACM0677 |
| 8 | AAZ3033, ACM0741, ACU4378 | 22 | ADI0213, AAH8773 |
| 9 | AAG1350, ACO1868 | 23 | ACJ5330, ADC9430 |
| 10 | AAU8556, ADN5159 | 24 | ACI7729, ACV3169 |
| 11 | AAG1277, ACA6737 | 25 | ADO9563, AAZ3033 |
| 12 | ABU5601, AAH8651 | 26 | ADA0629, ACP6154 |
| 13 | ABY9715, ACE4773 | 27 | AAH8868, ACQ9364 |
| 14 | ADH5015, ACE6023 | 28 | AAB6029, ABY9714 |

*Survey of BOLD Systems*. A download of all COI-5P from BOLD Systems cannot be done directly as the query is too large. Data was instead downloaded for each taxonomic category listed under Animals in BOLD Systems Taxonomy <http://boldsystems.org/index.php/TaxBrowser_Home>. Given the large size of Arthropoda dataset, the sub-taxa were downloaded separately and the same approach was adopted to Insecta dataset. The downloads were done either directly from the web browser or using BOLD Systems API, and “Combined data” was obtained for each. A separate download was also conducted for all data available under Data Releases section. As it is stated that this release only contains preliminary information of taxonomy, the updated metadata was obtained from the full download. Summary statistics available online in BOLD Systems were recorded and search terms were optimized based on the full data downloaded and these summary statistics. If the numbers were identical/very similar, those search terms were applied for 2015 dataset.

Table Sxxx. Metazoan data available in BOLD Systems Public database

|  | #Specimen records | # Seqs (2021 only) | # species (2021 only) | #COI-5P (2021/2015) | # BINs (2021/2015) |
| --- | --- | --- | --- | --- | --- |
| Total | **11,722,663** | **7,471,895** | **302,179** | **6,508,300/2,859,684** | **570,500/320,473** |
| Arthropoda | 10,321,635 | 6,572,565 | 230093 | 5,678,581/2,747,509 | 489,784/300,166 |
| Chordata | 842,287 | 518,022 | 35724 | 493,448/75,717 | 40,529/13,054 |
| Mollusca | 247,044 | 187,019 | 17764 | 179,263/10,096 | 20,919/2340 |
| Annelida | 104,937 | 61,922 | 4465 | 58,439/14,579 | 9272/3176 |
| Echinodermata | 57,578 | 29,768 | 1866 | 26,069/7684 | 2220/874 |
| Cnidaria | 29,900 | 20,353 | 3206 | 18,256/2004 | 1379/323 |
| Nematoda | 35,218 | 22,852 | 2609 | 14,490/657 | 1241/112 |
| Rotifera | 13,268 | 9307 | 509 | 9175/439 | 988/108 |
| Platyhelminthes | 38,963 | 29,271 | 2714 | 11,435/430 | 971/91 |
| Porifera | 7889 | 5099 | 1431 | 4537/71 | 730/44 |
| Nemertea | 5683 | 3858 | 414 | 3800/129 | 569/40 |
| Bryozoa | 4137 | 1842 | 245 | 1652/159 | 377/96 |
| Tardigrada | 3006 | 1769 | 232 | 1578/105 | 336/15 |
| Onychophora | 1393 | 1363 | 183 | 1353/5 | 246/2 |
| Gastrotricha | 1351 | 712 | 206 | 372/0 | 192/0 |
| Acanthocephala | 2306 | 2145 | 108 | 2032/2 | 182/2 |
| Chaetognatha | 1743 | 1344 | 102 | 1292/30 | 181/8 |
| Sipuncula | 1318 | 670 | 74 | 652/29 | 128/8 |
| Others (<100 BINs) | 3007 | 2014 | 234 | 1876/39 | 256/14 |
